# ntEmbd: Deep learning embedding for nucleotide sequences

**DOI:** 10.1101/2024.04.30.591806

**Authors:** Saber Hafezqorani, Ka Ming Nip, Inanc Birol

## Abstract

Enabled by the explosion of data and substantial increase in computational power, deep learning has transformed fields such as computer vision and natural language processing (NLP) and it has become a successful method to be applied to many transcriptomic analysis tasks. A core advantage of deep learning is its inherent capability to incorporate feature computation within the machine learning models. This results in a comprehensive and machine-readable representation of sequences, facilitating the downstream classification and clustering tasks. Compared to machine translation problems in NLP, feature embedding is particularly challenging for transcriptomic studies as the sequences are string of thousands of nucleotides in length, which make the long-term dependencies between features from different parts of the sequence even more difficult to capture. This highlights the need for nucleotide sequence embedding methods that are capable of learning input sequence features implicitly. Here we introduce ntEmbd, a deep learning embedding tool that captures dependencies between different features of the sequences and learns a latent representation for given nucleotide sequences. We further provide two sample use cases, describing how learned RNA features can be used in downstream analysis. The first use case demonstrates ntEmbd*’*s utility in classifying coding and noncoding RNA benchmarked against existing tools, and the second one explores the utility of learned representations in identifying adapter sequences in nanopore RNA-seq reads. The tool as well as the trained models are freely available on GitHub at https://github.com/bcgsc/ntEmbd

## INTRODUCTION

Recent advancements in high-throughput sequencing technologies have enabled us to generate a comprehensive global view of the transcriptome for various species and cell types, allowing for the cataloging of transcript species, determining transcriptional structures, and quantifying transcript expression levels, which are all crucial for studying disease mechanisms [1–5]. This necessitated the development of computational methods to effectively capture and analyze RNA sequences. Machine learning algorithms, with their capability to handle large and complex datasets, can be particularly useful in tackling this problem.

The effectiveness of traditional machine learning algorithms is largely influenced by the data representation they employ, that is, on how each feature is computed. The quality and relevance of these features significantly impact classification model*’*s outcomes as human-designed features may introduce biases into model predictions. Therefore, identifying optimal features is challenging for numerous bioinformatics classification tasks. A promising solution to this challenge is representation learning where the model learns to identify the most effective feature set implicitly and summarize them in vectors of real number, named embeddings.

Traditional embedding approaches, such as one-hot encoding or *k*-mer based representations, are limited in their ability to capture the contextual dependencies inherent in nucleotide sequences. These methods often result in high-dimensional, sparse data representations that are not conducive to efficient computation or effective learning. Moreover, while Word2Vec [6] and FastText [7–9], which borrow from successes in natural language processing, provide a more sophisticated approach compared to one-hot encoding by considering local context through word or subword embeddings, they do not fully appreciate the sequential order of nucleotides or long-range interactions relevant for RNA structure and function. Therefore, methods based on these approaches are limited in capturing the full complexity of sequence-dependent features since they assign the same vector to the same word, regardless of its context [10–14].

Embedding methods reduce nucleotide sequences to compact, fixed-length vector representations while effectively learning their features. These features can be applied on different downstream tasks where the traditional approaches often depend on explicit, handcrafted features, which can introduce biases and may fail to encapsulate the full spectrum of sequence intricacies [15–18]. Such engineered features, while beneficial in certain contexts, are derived from prior biological knowledge and assumptions that limit the model*’*s ability build upon those features to reveal novel insights and to generalize across different RNA sequence landscapes. Moreover, these manual features might overlook subtle, yet biologically important patterns within the nucleotide sequences, due to their reliance on what is already understood rather than what can be discovered through data-driven approaches.

Deep learning offers a promising avenue to address these challenges in transcriptomic studies. An advantage of deep learning is its capacity to integrate feature extraction directly within machine learning algorithms. This integration yields representations of sequences that are both rich in information and readily interpretable by machines, thereby streamlining the creation of downstream models for classification and clustering. In the protein domain, for instance, deep learning techniques have been leveraged to learn representations from amino acid sequences [19,20]. On the other hand, feature embedding is particularly challenging for transcriptomic studies as the sequences are much longer and its alphabet consist of four letters, which makes it more difficult to capture the long-term dependencies between features from different parts of the sequence. This highlights the need for nucleotide sequence embedding methods that are tailored to this problem domain and are capable of learning input sequence features implicitly.

Here, we present ntEmbd, an embedding method to learn a fixed-dimensional representation for nucleotide sequences using deep learning autoencoder architecture. We demonstrate use case of ntEmbd embeddings in two downstream analysis: 1) RNA coding potential assessment task classifying coding versus noncoding sequences, and 2) Detecting adapter sequences in long reads generated by Oxford Nanopore Technologies (ONT, Oxford, UK).

## RESULTS

### Application in coding and noncoding classification

We first assessed the effectiveness of the ntEmbd learned representations in a functional annotation task involving the classification of RNA sequences into coding or noncoding transcripts. We selected the RNA sequences from the human reference transcriptome and labeled them as either protein-coding (mRNA) or noncoding (ncRNA) according to the biotype classes. We filtered these sequences based on three different maximum length thresholds (1500, 2000, and 3000 nt) and pre-padded them to get unified lengths. This resulted in 14,090, 26,488, and 51,058 mRNA sequences and 10,728, 13,089, and 15,937 ncRNA sequences respectively. We trained ntEmbd models using these sequences and generated the learned representation for each RNA sequence. Utilizing the ntEmbd-generated embeddings along with respective sequence labels, we then trained supervised classifiers to distinguish coding and noncoding RNAs. The ensemble model achieved an accuracy of 0.874, a sensitivity of 0.957, receiving an F1 score of 0.920. The model*’*s specificity decreased as the length of input sequences increased, whereas both sensitivity and precision increased, resulting in higher overall F1 scores with longer input sequences.

### Benchmarking against other RNA coding potential assessment tools

We further assessed the effectiveness of ntEmbd generated embeddings in RNA coding potential assessment task by comparing its performance against five other existing state-of-the-art predictors in this domain [17,21–24]. Table 1 summarizes the comparison of classification performance using trained models on the mRNN dataset [24]. The training set consists of 87,188 mRNA and 24,421 lncRNA sequences and the test set includes an equal number of mRNA and lncRNA sequences, each with 500 sequences, designed to test classification capabilities under stringent conditions with mRNAs with short ORFs (≤50 codons in the GENCODE annotation) and lncRNAs with long untranslated ORFs (≥50 codons). The classifier built upon ntEmbd embeddings outperformed other tools in almost every metric, notably achieving high accuracy (0.884), recall (0.938), and F1-score (0.886), only falling short on precision (0.840).

**Table 1:** Benchmarking ntEmbd embedding in RNA coding potential assessment task.

| Tools | Sensitivity | Precision | F1 score | Accuracy |
| --- | --- | --- | --- | --- |
| **RNASamba** | 0.774 | 0.917 | 0.839 | 0.852 |
| **CPAT** | 0.528 | **0.967** | 0.683 | 0.755 |
| **CPC2** | 0.384 | 0.932 | 0.543 | 0.678 |
| **FeeLnc** | 0.598 | 0.937 | 0.730 | 0.779 |
| **mRNN** | 0.798 | 0.936 | 0.861 | 0.872 |
| **ntEmbd** | **0.938** | 0.840 | **0.886** | **0.884** |

### Impact of sequence completeness on RNA classification accuracy

We also explored the impact of RNA sequence completeness on the performance of classifiers in distinguishing coding from noncoding transcripts. We first generated three set of sequences from complete RNA transcripts: those containing the longest open reading frame of the RNA transcript (referred as full set in this analysis), those containing 5*’*, 3*’*, or both 5*’* and 3*’* truncated sequences (referred as partial set), and lastly a combination of both. Next, we extracted embeddings for these full and partial-length transcripts using a pre-trained ntEmbd model and assessed the distance between these embeddings. Visualizing the UMAP plot of full and partial-length transcripts*’* embedding vectors using the Tensorflow Embedding Projector [25] indicated a noticeable separation between the two classes (Figure 1).

**Figure 1:**
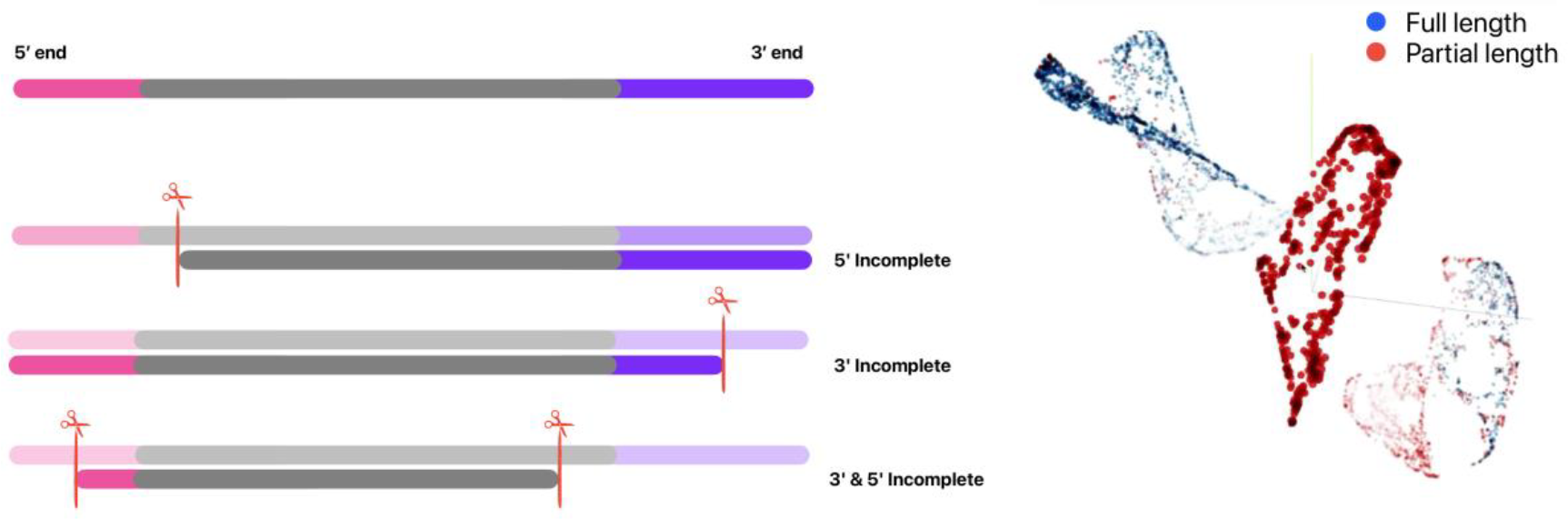
Visualizing full and partial-length transcripts*’* embedding vectors

Next, we trained three classifiers using these embeddings as input: one model with only full-length, another with only partial-length, and a third with a combination of both sequence types. We also employed the pre-trained ntEmbd model to get the embedding vectors of the original RNA transcript sequences studied in this analysis and used them as test set to evaluate the above-mentioned three models. The model trained on both sequence types (full + partial) demonstrated higher accuracy, with a score of 0.646, in comparison to the models trained exclusively on full-length or partial-length sequences, which scored 0.564 and 0.589, respectively (Table 2). This suggests that training on a dataset that mirrors the variation found in actual RNA sequences— comprising both complete and incomplete transcripts—enhances the classifier*’*s ability to accurately predict RNA coding potential in real-world scenarios.

**Table 2:** Coding and noncoding RNA classification metrics studying the effect of sequence completeness.

| Model | Sensitivity | Precision | F1 score | Accuracy |
| --- | --- | --- | --- | --- |
| **Only Full** | **0.996** | 0.559 | 0.716 | 0.564 |
| **Only Partial** | 0.974 | 0.576 | 0.724 | 0.589 |
| **Both** | 0.859 | **0.632** | **0.728** | **0.646** |

### Transfer learning with ntEmbd for identifying ONT adapters

Adapters are essential component of the ONT sequencing workflows, and their sequences are commonly found in the final output reads. These sequences are typically either trimmed during base calling (in which strand-specificity is lost) or in post-processing analysis, making final sequences ready for downstream tasks. We investigated the utility of ntEmbd embeddings for detecting adapter sequences in ONT sequencing data through a transfer learning approach. First, we processed ONT cDNA reads from a mouse replicate (ENCFF232YSU) of the LRGASP project [26] with pychopper [27] to identify ones with and without primer sequences. We used a pre-trained ntEmbd model to learn features of these sequences and generated their embedding vectors. Next, we trained two classifiers to distinguish reads containing primers from those without. For the first classifier, we used the ntEmbd embeddings as input whereas for the second one, we fed *K*-mer embeddings for training. The latter is a collection of unique integer values assigned to each 3-mer within the RNA sequence.

The classifier utilizing ntEmbd embeddings achieved an accuracy of 0.91, with both sensitivity and precision exceeding those achieved by a classifier using *K*-mer embeddings (0.977 and 0.862 compared to 0.951 and 0.690) as demonstrated in Table 3. This analysis, conducted on a model not originally trained on ONT reads, showcases ntEmbd*’*s transfer learning capability for feature learning and suggests its potential for broader applications within nanopore sequencing workflows.

**Table 3:**
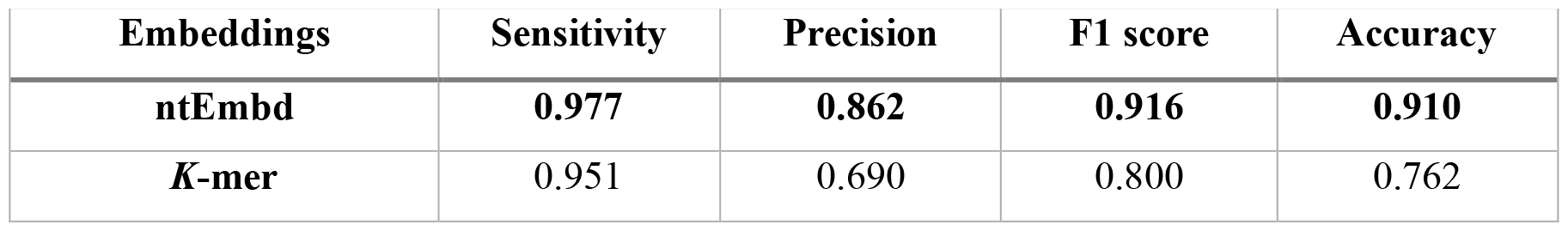
Transfer learning with ntEmbd for identifying ONT adapters.

## METHODS

### Architecture of ntEmbd

ntEmbd*’*s general workflow for encoding RNA sequences into embeddings is summarized in Figure 2. It employs an unsupervised Bi-LSTM autoencoder architecture [28,29] to process high-dimensional nucleotide sequences into a compact latent representation. Initially, the input nucleotide sequences are preprocessed using an encoding scheme that is generalized to account for the IUPAC nucleotide wildcards [30] (Figure 3). These sequences are then fed to the autoencoder component, which consists of an encoder and a decoder. The encoder segment of the autoencoder then learns to map the data, denoted by *x*, into a reduced-dimensional latent space, z. This transformation is pivotal for capturing the essential features of the sequences without direct supervision. Subsequently, the decoder reverses this process, mapping from the latent space back to the data space to generate a reconstruction of the original sequences, denoted by *y*.

**Figure 2:**
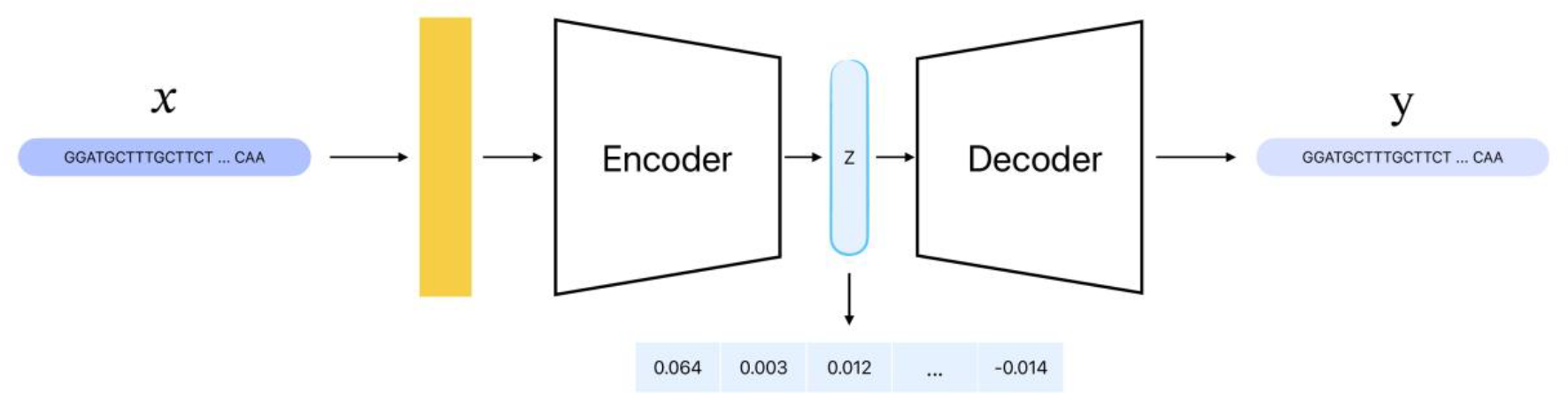
Overall view of ntEmbd architecture

**Figure 3:**
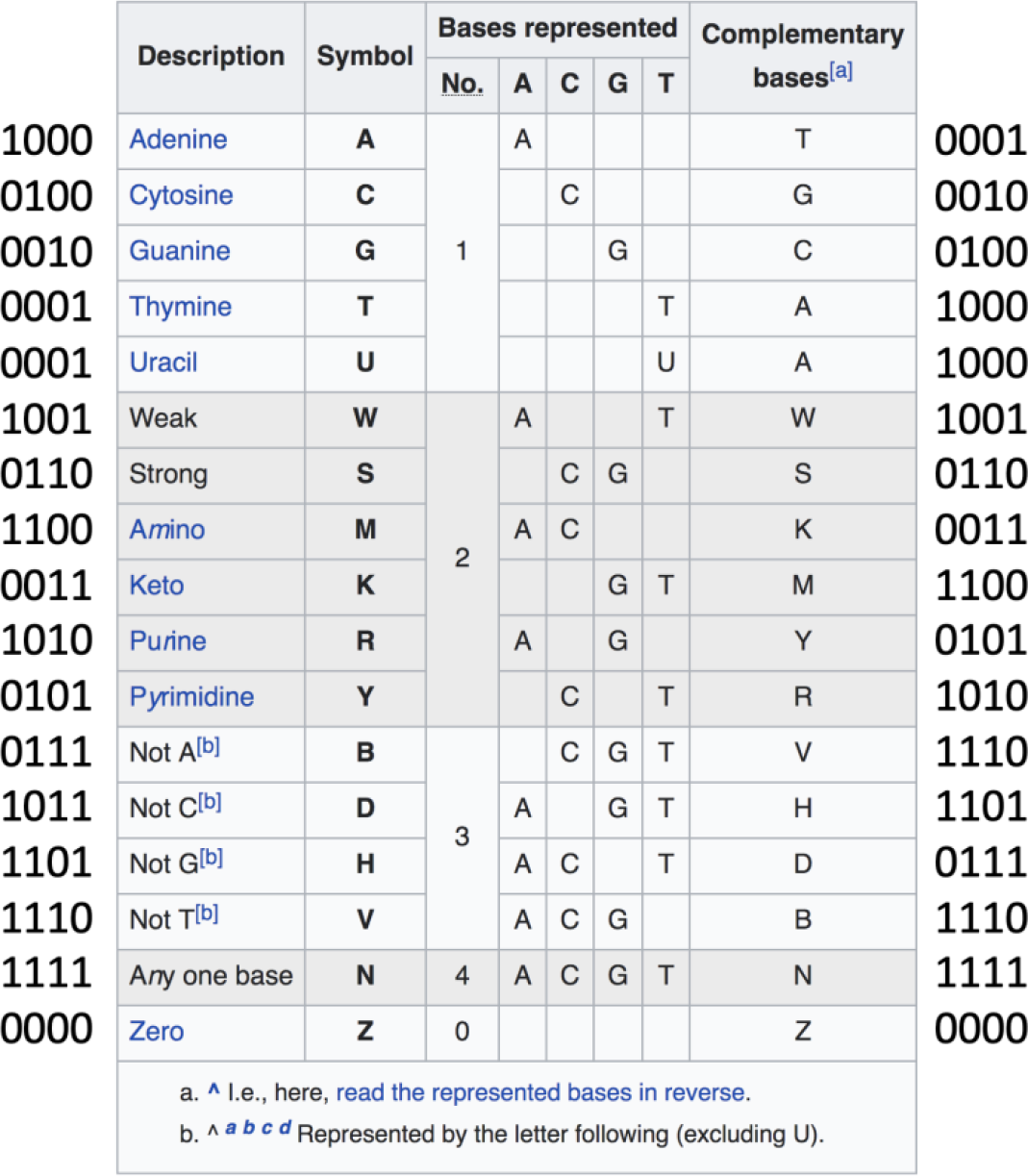
IUPAC degenerate base symbols. The illustration is adopted from Nucleic acid notation Wikipedia page [31].

The model*’*s training objective is to minimize the reconstruction loss, *L*(*x, y*), which measures the distance of the reconstituted sequences to the original ones. Through this process, the encoder is encouraged to retain only the most important features in the latent space representation. These features are encoded as real-numbered vectors in z.

For the above-mentioned loss function calculation to work, the implementation should satisfy the requirements for a distance measure between sequences of the same length to be a metric, meaning that the distance to self is zero, the distance between two vectors is symmetric, and finally it follows the triangle inequality. To quantify the loss function and measure the distance between vector representation of input sequences and their reconstructed observation, the model incorporates the following formula where for vectors *x* and *y*, the angular distance is calculated using the arccosine of their dot product, scaled by a factor of 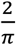 to ensure the loss ranges between 0 and 1.

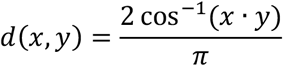

This workflow design allows ntEmbd to identify and represent the biophysical properties of RNA sequences in the latent space as vectors of real numbers, without explicitly defining these properties in prior. The resulting features from the latent space are then utilized in various downstream tasks as demonstrated in two use cases described in the Results section.

### Data preparation and training strategy

To train ntEmbd, we utilized full-length reference human transcript sequences. In the pre-processing stage, sequences were first adjusted to a uniform length through padding. ntEmbd also provides the functionality for users to truncate sequences based on minimum and maximum length parameters. This length standardization facilitates batch processing, which is critical for the computational efficiency of the training process required by modern deep learning frameworks such as TensorFlow and Pytorch [32,33]. Following this, the dataset was prepared for validation and testing: 20% of the data was reserved for final evaluation, while the remaining 80% was used for training. Within this training subset, a further split was executed to form a validation set which is used for monitoring the model training performance. This split ratio is 75/25 percent by default and users have the option to set their own preferred values when training a ntEmbd model.

ntEmbd employs early stopping approach to prevent overfitting during training. It monitors the performance of the model on validation set with a *patience* variable set to 10 by default. This specifies the number of epochs it waits after the last time the model*’*s performance improved on the validation set. We also incorporated a model checkpoint callback that saves the best model in each step.

### Classification models

Following the generation of learned representations for each RNA sequence using the ntEmbd model, these embedding vectors serve as inputs for training several classifiers to facilitate classification analysis. The models trained include Multilayer Perceptron (MLP), K-Nearest Neighbors (KNN), Gradient Boosting (GB), and Random Forest. To optimize performance, an ensemble method is employed, wherein the best-performing model from these classifiers is selected based on their collective accuracy and used in downstream classification tasks.

### Hyperparameter tuning

Hyperparameters optimization is a critical step in enhancing the model*’*s performance. For this purpose, the Optuna package, a hyperparameter optimization framework, was employed [34]. Optuna offers a versatile and user-friendly interface for hyperparameter tuning, which can be executed in two modes in ntEmbd pipeline: a standalone mode dedicated solely to hyperparameter tuning and an integrated mode that allows for hyperparameter tuning concurrently with model training. The goal of an Optuna study is to find out the optimal set of hyperparameter values through multiple trials. Users have the flexibility to select from a variety of `Samplers`, which are algorithms to suggest parameter values, and `Pruners`, which are methods to stop unpromising trials early, thus improving computational efficiency.

We implemented an automated search space suggestion feature within ntEmbd that analyzes a subset of the training data before the commencement of tuning and recommends an appropriate search space for the hyperparameters, thus streamlining the tuning process by narrowing down the hyperparameter search space. This component analyzes the size of training data as well as the length distribution of input sequences and suggests a value range for autoencoder latent space size, number of LSTM units, and batch sizes for the subsequent Optuna hyperparameter tuning process.

ntEmbd employs a 5-fold cross-validation approach where it initializes an Optuna study and records the best loss and parameters for each fold. Finally, it aggregates the best hyperparameters across folds using voting strategy for categorical parameters and averaging for continuous parameters. These aggregated best parameters are then used in training the ntEmbd autoencoder model or reported to the user in the case of running hyperparameter tuning module on standalone mode.

The Optuna run within ntEmbd also supports parallelization, allowing multiple trials to be conducted simultaneously. This is facilitated using a rational database (RDB) where the information of each trial is written into RDB instead of a memory. This approach not only expedites the tuning process but also ensures the robustness and reproducibility of the optimization. Additionally, ntEmbd incorporates Optuna*’*s visualization capabilities to aid users in interpreting the optimization process and making informed decisions about parameter selection.

### Modular design

The ntEmbd pipeline is designed with a modular structure, comprising discrete components for training, embedding, visualization, and analysis. This means that researchers can use the pipeline to train their own model, get the embedding vectors for given RNA sequences using a pre-trained model, visualize the embedding vectors, or perform only hyperparameter tuning analysis.

## CONCLUSION

We introduced ntEmbd, a deep learning embedding technique to learn feature representation of nucleotide sequences and demonstrated the power of ntEmbd-generated representations in two use cases. First, we assessed the effectiveness of ntEmbd generated embeddings in classifying coding versus noncoding RNA transcripts by benchmarking its performance against five other tools. We also studied the impact of RNA sequence completeness in model*’*s classification performance. We further applied the ntEmbd embeddings on Nanopore adapter sequence detection task and demonstrated its transfer learning capability for feature learning. We believe that ntEmbd has the potential to be a valuable tool for other sequence classification and clustering tasks when fine-tuned to the specific problem space.

## DECLARATIONS

### Data and software availability

ntEmbd is implemented in Python. The source code and pre-trained models used in this study are available on Github: https://github.com/bcgsc/ntEmbd

### Authors’ contributions

SH and IB conceived the study. SH designed and developed ntEmbd under the supervision of IB. SH conducted the benchmarking experiments. KMN aided with experimental designs. SH took the lead in writing the manuscript with input from all authors.

## References

1. Neve J, Patel R, Wang Z, Louey A, Furger AM. Cleavage and polyadenylation: Ending the message expands gene regulation. RNA Biol. 2017 Jul 3;14(7):865–90.

2. Wang Z, Gerstein M, Snyder M. RNA-Seq: a revolutionary tool for transcriptomics. Nature Reviews Genetics 2008 10:1 [Internet]. 2009 Jan [cited 2024 Feb 4];10(1):57–63. Available from: https://www.nature.com/articles/nrg2484

3. Costa V, Aprile M, Esposito R, Ciccodicola A. RNA-Seq and human complex diseases: recent accomplishments and future perspectives. European Journal of Human Genetics. 2013;21(2):134–42.

4. Prokop JW, Shankar R, Gupta R, Leimanis ML, Nedveck D, Uhl K, et al. Virus-induced genetics revealed by multidimensional precision medicine transcriptional workflow applicable to COVID-19. Physiol Genomics. 2020 Jun 1;52(6):255–68.

5. Mortazavi A, Williams BA, McCue K, Schaeffer L, Wold B. Mapping and quantifying mammalian transcriptomes by RNA-Seq. Nature Methods 2008 5:7. 2008 May 30;5(7):621–8.

6. Mikolov T, Chen K, Corrado G, Dean J. Efficient Estimation of Word Representations in Vector Space. 1st International Conference on Learning Representations, ICLR 2013 - Workshop Track Proceedings. 2013 Jan 16;

7. Bojanowski P, Grave E, Joulin A, Mikolov T. Enriching Word Vectors with Subword Information. Trans Assoc Comput Linguist. 2017 Dec 1;5:135–46.

8. Joulin A, Grave E, Bojanowski P, Douze M, Jégou H, Mikolov T. FastText.zip: Compressing text classification models. 2016 Dec 12;

9. Joulin A, Grave E, Bojanowski P, Mikolov T. Bag of Tricks for Efficient Text Classification. 15th Conference of the European Chapter of the Association for Computational Linguistics, EACL 2017 - Proceedings of Conference. 2016 Jul 6;2:427– 31.

10. Ng P. dna2vec: Consistent vector representations of variable-length k-mers. 2017 Jan 23;

11. Dai H, Umarov R, Kuwahara H, Li Y, Song L, Gao X. Sequence2Vec: a novel embedding approach for modeling transcription factor binding affinity landscape. Bioinformatics. 2017 Nov 15;33(22):3575–83.

12. Asgari E, Mofrad MRK. Continuous Distributed Representation of Biological Sequences for Deep Proteomics and Genomics. PLoS One. 2015 Nov 10;10(11):e0141287.

13. Zou Q, Xing P, Wei L, Liu B. Gene2vec: gene subsequence embedding for prediction of mammalian N6-methyladenosine sites from mRNA. RNA. 2019 Feb 1;25(2):205–18.

14. Menegaux R, Vert JP. Continuous Embeddings of DNA Sequencing Reads and Application to Metagenomics. https://home.liebertpub.com/cmb. 2019 Jun 6;26(6):509– 18.

15. Tong X, Liu S. CPPred: coding potential prediction based on the global description of RNA sequence. Nucleic Acids Res. 2019 May 7;47(8):e43–e43.

16. Hu L, Xu Z, Hu B, Lu ZJ. COME: a robust coding potential calculation tool for lncRNA identification and characterization based on multiple features. Nucleic Acids Res. 2017 Jan 9;45(1):e2–e2.

17. Kang YJ, Yang DC, Kong L, Hou M, Meng YQ, Wei L, et al. CPC2: a fast and accurate coding potential calculator based on sequence intrinsic features. Nucleic Acids Res. 2017 Jul 3;45(W1):W12–6.

18. Kong L, Zhang Y, Ye ZQ, Liu XQ, Zhao SQ, Wei L, et al. CPC: assess the protein-coding potential of transcripts using sequence features and support vector machine. Nucleic Acids Res. 2007 Jul 1;35(Suppl_2):W345–9.

19. Heinzinger M, Elnaggar A, Wang Y, Dallago C, Nechaev D, Matthes F, et al. Modeling aspects of the language of life through transfer-learning protein sequences. BMC Bioinformatics. 2019 Dec 17;20(1):1–17.

20. Elnaggar A, Heinzinger M, Dallago C, Rehawi G, Wang Y, Jones L, et al. ProtTrans: Toward Understanding the Language of Life Through Self-Supervised Learning. IEEE Trans Pattern Anal Mach Intell. 2022 Oct 1;44(10):7112–27.

21. Wucher V, Legeai F, Hédan B, Rizk G, Lagoutte L, Leeb T, et al. FEELnc: a tool for long non-coding RNA annotation and its application to the dog transcriptome. Nucleic Acids Res. 2017 May 5;45(8):e57–e57.

22. Wang L, Park HJ, Dasari S, Wang S, Kocher JP, Li W. CPAT: Coding-Potential Assessment Tool using an alignment-free logistic regression model. Nucleic Acids Res. 2013 Apr 1;41(6):e74–e74.

23. Camargo AP, Sourkov V, Pereira GAG, Carazzolle MF. RNAsamba: neural network-based assessment of the protein-coding potential of RNA sequences. NAR Genom Bioinform. 2020 Mar 1;2(1).

24. Hill ST, Kuintzle R, Teegarden A, Merrill E, Danaee P, Hendrix DA. A deep recurrent neural network discovers complex biological rules to decipher RNA protein-coding potential. Nucleic Acids Res. 2018 Sep 19;46(16):8105–13.

25. Smilkov D, Brain G, Thorat N, Nicholson C, Reif E, Viégas FB, et al. Embedding Projector: Interactive Visualization and Interpretation of Embeddings. 2016 Nov 16;

26. Pardo-Palacios FJ, Wang D, Reese F, Diekhans M, Carbonell-Sala S, Williams B, et al. Systematic assessment of long-read RNA-seq methods for transcript identification and quantification. bioRxiv. 2023 Jul 27;3:2023.07.25.550582.

27. epi2me-labs/pychopper: cDNA read preprocessing - https://github.com/epi2me-labs/pychopper.

28. Hochreiter S, Schmidhuber J. Long Short-Term Memory. Neural Comput. 1997 Nov 15;9(8):1735–80.

29. Schuster M, Paliwal KK. Bidirectional recurrent neural networks. IEEE Transactions on Signal Processing. 1997;45(11):2673–81.

30. Cornish-Bowden A. Nomenclature for incompletely specified bases in nucleic acid sequences: recommendations 1984. Nucleic Acids Res. 1985 May 5;13(9):3021.

31. Nucleic acid notation - Wikipedia - https://en.wikipedia.org/wiki/Nucleic_acid_notation.

32. Abadi M, Agarwal A, Barham P, Brevdo E, Chen Z, Citro C, et al. TensorFlow: Large-Scale Machine Learning on Heterogeneous Distributed Systems. 2016 Mar 14;

33. Paszke A, Gross S, Massa F, Lerer A, Bradbury J, Chanan G, et al. PyTorch: An Imperative Style, High-Performance Deep Learning Library. Adv Neural Inf Process Syst. 2019 Dec 3;32.

34. Akiba T, Sano S, Yanase T, Ohta T, Koyama M. Optuna: A Next-generation Hyperparameter Optimization Framework. Proceedings of the ACM SIGKDD International Conference on Knowledge Discovery and Data Mining. 2019 Jul 25;2623– 31.

